# Genome-Guided Discovery and Biosynthetic Characterization of Burkstatins, Bicyclic Microviridins with Protease Inhibitory Activity

**DOI:** 10.64898/2026.11.12.766270

**Authors:** Zihao Cheng, Allison S. Walker

## Abstract

Microviridins are ribosomally synthesized and post-translationally modified peptides characterized by a highly constrained macrocyclic scaffold and potent serine protease inhibitory activity. Unlike canonical tricyclic microviridins, bicyclic microviridins, exemplified by marinostatins, lack the characteristic ω-amide linkage. The biosynthetic diversity and biological activities of bicyclic microviridins remain poorly explored. Here, genome mining and phylogenetic analyses revealed diverse lineages of putative bicyclic microviridins, including a conserved family of multicore precursors in Burkholderiaceae. We characterized a representative biosynthetic gene cluster from *Paraburkholderia* sp. CNPSo 3076 and demonstrated that a single ATP-grasp ligase, BksD, catalyzes the formation of two ω-ester macrocycles within each of the two precursor core regions. Heterologous reconstitution and tandem mass spectrometric analyses enabled the production and structural characterization of two bicyclic peptides, designated burkstatin 1 and burkstatin 2. Despite sharing a conserved bicyclic scaffold, the two peptides exhibited distinct serine protease inhibitory profiles. Burkstatin 2 potently inhibited subtilisin and elastase, whereas burkstatin 1 displayed weaker subtilisin inhibition and no detectable elastase inhibition. These findings expand the known biosynthetic and functional diversity of bicyclic microviridins, demonstrate the capacity of a single ATP-grasp ligase to modify multiple core peptides and highlight the influence of core sequence variation on protease inhibitory activity.

## INTRODUCTION

Ribosomally synthesized and post-translationally modified peptides (RiPPs) constitute a subset of natural products with diverse bioactivities.^1–3^ RiPP biosynthetic gene clusters (BGCs) generally encode a precursor peptide, modifying enzymes responsible for its post-translational modification, proteases, transporters and other accessory proteins.^1^ The precursor peptide generally comprises a leader peptide (LP) region that facilitates recognition by biosynthetic enzymes and a core peptide (CP) region that undergoes diverse modifications, including methylation, oxidation, dehydration, and covalent crosslinking.^1^ Following modification, the mature core peptide is typically released from leader peptide through proteolytic cleavage.^1^ Among the diverse post-translational modifications found in RiPP biosynthesis, many lead to the formation of macrocycles, exemplified by families of RiPPs such as lanthipeptides^4^, graspetides^5^, and lasso peptides.^6^ Compared to linear peptides, macrocyclic peptides are more conformationally constrained through covalent crosslinks between residues, often resulting in higher resistance to proteolysis and better potency against targets.^7,8^ Macrocyclic RiPPs therefore hold considerable potential for drug discovery by providing biosynthetic access to structurally complex, conformationally constrained peptide scaffolds with favorable pharmacological properties that are often challenging to synthesize through conventional chemical synthesis.^9^

Microviridins are a family of macrocyclic RiPPs initially discovered from cyanobacteria and are characterized as potent and selective inhibitors of serine proteases.^5^ Canonical microviridin precursor peptides contain a conserved heptapeptide motif, PFFARFL^10^, in the leader peptide that facilitates ATP-grasp ligase recognition together with a highly conserved core sequence commonly represented as TxKxPSDx(D/E)(D/E).^11–13^ Macrocyclization is catalyzed sequentially by two homologous ATP-grasp ligases. In microviridin biosynthesis, MvdD first installs two ω-ester linkages between the side-chain hydroxyl groups of Thr and Ser and carboxylate groups of Asp/Glu, whereas MvdC subsequently forms an ω-amide linkage between Lys and a C-terminal Asp/Glu residue.^11^ The resulting two lactones and one lactam generate the characteristic cage-like topology of mature microviridins (Figure 1).^11^ Subsequent studies uncovered diverse precursor peptides distinct from canonical microviridins that are similarly macrocyclized by ATP-grasp ligases through side-chain ester or amide linkages, establishing the broader RiPP family now known as graspetides.^12,14^

**Figure 1.**
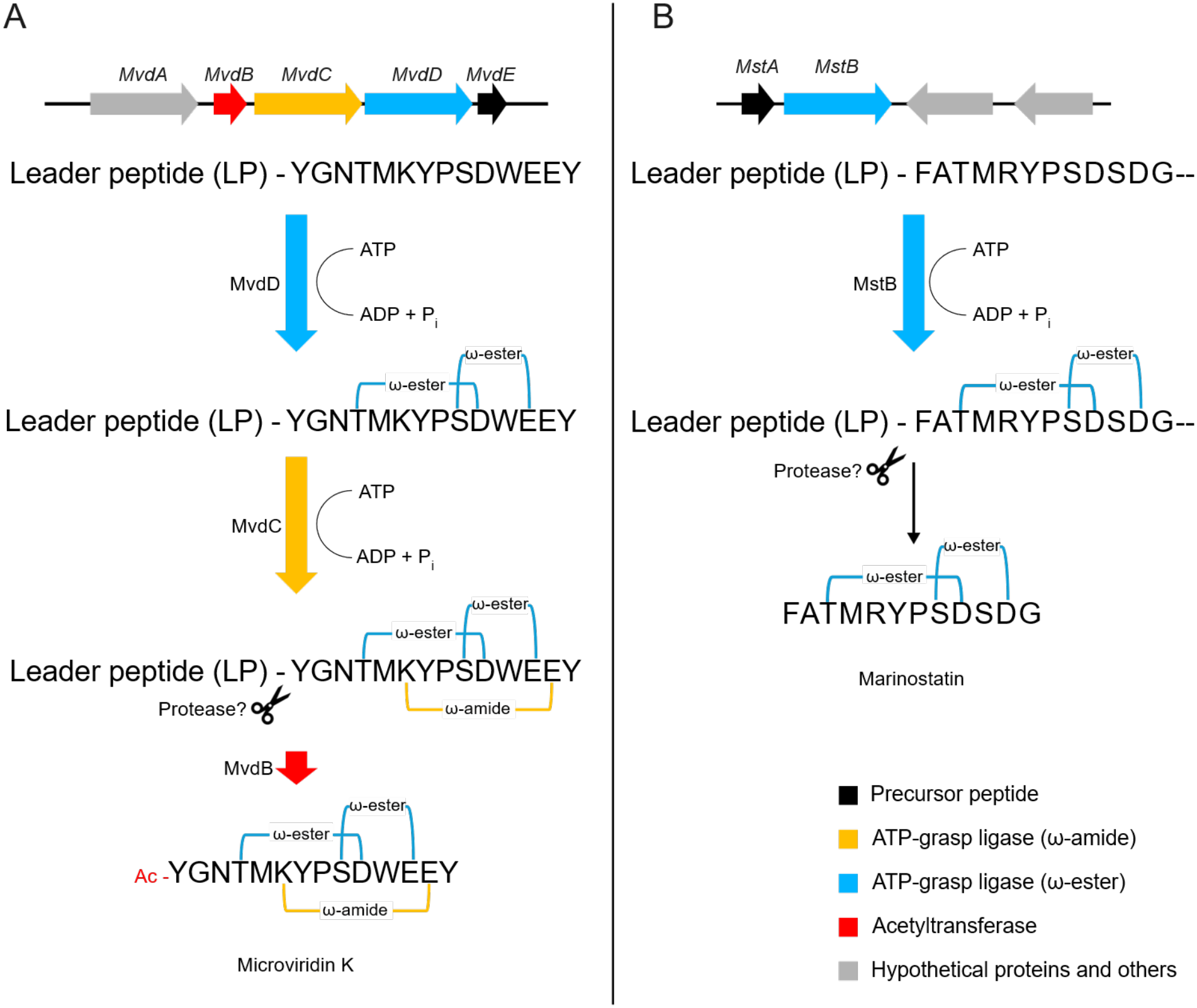
Comparison of BGC and macrocyclization pathways of microviridin K (left) and marinostatin (right).

In contrast to the residues required for ring formation, several positions presented within or adjacent to the constrained microviridin scaffold tolerate considerable sequence variation.^15–17^ This division between a conserved structural framework and variable substrate-interacting residues provides a molecular basis for diversification of protease specificity. Mutagenesis of variable core positions, has generated microviridin libraries with altered protease selectivity.^15,18^ In particular, the residue between T and K of mature microviridins is a major determinant of inhibitory selectivity.^15^ Natural sequence comparisons, structural studies, and mutational analyses have shown that substitution at this position can redirect inhibition among trypsin-, chymotrypsin-, elastase-, and subtilisin-like proteases. For example, introduction of basic residues favors trypsin inhibition, whereas hydrophobic residues can confer potent activity toward elastase or other serine proteases.^15,18^ Consequently, the highly constrained microviridin macrocycles provide a proteolytically resistant framework while retaining a genetically tunable recognition residues that can alter target preference.

An additional layer of diversity arises from microviridin precursors containing multiple core motifs within a single polypeptide. Genome-mining studies have identified microviridin BGCs containing precursors with as many as five core peptides, with multicore architectures distributed across multiple bacterial phyla.^19^ Biochemical characterization of the three-core AMdnA precursor from *Anabaena* sp. PCC 7120 demonstrated that its cognate ATP-grasp ligase distributively modifies the individual cores, with modification of one core having little apparent influence on macrocyclization of the others.^16^ Notably, the three cores are not processed identically, as the first core of the predominant modified precursor receives only one macrolactone whereas the remaining cores receive two.^16^ Despite the prevalence of this architecture, the biological significance and downstream maturation of multicore microviridins remain poorly understood. Conserved intercore motifs can provide potential proteolytic cleavage sites, yet individual mature microviridins were not isolated from multicore systems in previous studies.^20^ More broadly, microviridin maturation requires proteolytic removal of precursor sequences following macrocyclization, but dedicated maturation proteases are not universally apparent within their BGCs.^19,21,22^ Thus, in pathways lacking dedicated proteolytic machinery, processing by endogenous host proteases could provide an additional layer of diversification, potentially releasing products that share a conserved macrocyclic core while differing in their N- or C-terminal residues.^21^ How multicore architecture contributes to the production and diversification of mature microviridins therefore remains an unresolved aspect of their biosynthesis.^20^

Marinostatins represent a structurally simplified branch of the microviridin family.^23–26^ Like microviridins, marinostatins are serine protease inhibitors containing two ω-ester-linked macrocycles; however, their core sequences lack the conserved Lys required for formation of the third, ω-amide-linked ring.^23^ Accordingly, characterized marinostatin biosynthetic loci encode an ester-forming ATP-grasp ligase related to MvdD but lack the MvdC-type ligase required for lactam formation (Figure 1).^21^ Conservation of the microviridin-like leader peptide and lactone-forming machinery, together with phylogenetic analyses of the associated biosynthetic proteins, has led to the proposal that marinostatins represent a derived bicyclic lineage of microviridins that arose through loss of the lactam-forming ATP-grasp ligase.^5^ While multicore architectures are widespread among predicted microviridin precursors, the biosynthesis and maturation of multicore marinostatin-like precursors remain largely unexplored.

This evolutionary relationship raises broader questions regarding how frequently the canonical tricyclic microviridin architecture has transitioned to bicyclic forms, how these architectures are distributed across bacterial lineages, and whether loss of the third macrocycle alters the functional properties of the resulting peptides. Recent genome-mining studies have greatly expanded the recognized taxonomic and structural diversity of microviridin-like graspetides, identifying biosynthetic loci across Cyanobacteriota, Bacteroidota, and Pseudomonadota and revealing multicore precursors and noncanonical combinations of ATP-grasp ligases.^11,12,19^ However, the biosynthetic and functional diversity of marinostatin-like, bicyclic microviridins remains comparatively underexplored.

Here, we used genome mining and precursor-centered phylogenetic analysis to investigate the diversity and evolution of bicyclic microviridins. We identify multiple phylogenetically distinct lineages of marinostatin-like precursors, including multicore architectures and biosynthetic loci with unusual ATP-grasp ligase complements, revealing a dynamic relationship between precursor architecture and the capacity for ω-amide cyclization. In particular, we uncover a conserved lineage of multicore bicyclic microviridins in *Paraburkholderia*, a genus from which marinostatin-like natural products have not, to our knowledge, been experimentally characterized. Guided by this analysis, we selected a representative biosynthetic gene cluster from *Paraburkholderia* sp. CNPSo 3076 and investigated the biosynthesis, structures, and protease inhibitory activities of its encoded bicyclic microviridins.

## RESULTS AND DISCUSSION

### Phylogenetic Analysis Reveals Diverse Lineages of Bicyclic Microviridins

To investigate the biosynthetic diversity of bicyclic microviridins, we surveyed microviridin precursor peptides identified through antiSMASH 5^27^ genome mining across taxonomically diverse bacteria, including members of Cyanobacteriota, Bacteroidota, and Pseudomonadota. In total, we identified 36 putative marinostatin-like biosynthetic gene clusters (BGCs) containing precursor peptides with Lys-deficient core sequences that lack the conserved lysine required for formation of the characteristic microviridin macrolactam (SI, Table S3). Because these core sequences resemble the bicyclic architecture of marinostatins, we classified them as putative bicyclic microviridin precursors and examined their evolutionary relationships in greater detail.

We extracted the marinostatin-like precursor sequences and constructed a midpoint-rooted maximum-likelihood phylogenetic tree (Figure 2). The resulting tree revealed substantial sequence and architectural diversity among putative bicyclic microviridins. Although some precursors contained a single Lys-deficient core, numerous sequences encoded two or more predicted core regions within a single precursor. Several multicore precursors were particularly notable because they combined Lys-deficient marinostatin-like cores with Lys-containing microviridin-like cores. Moreover, bicyclic core-containing precursors were distributed across phylogenetically separated branches and distantly related bacterial taxa rather than forming a single monophyletic lineage. Together, these patterns suggest that the Lys-dependent ω-amide cyclization has been gained or lost independently multiple times across different lineages.

**Figure 2.**
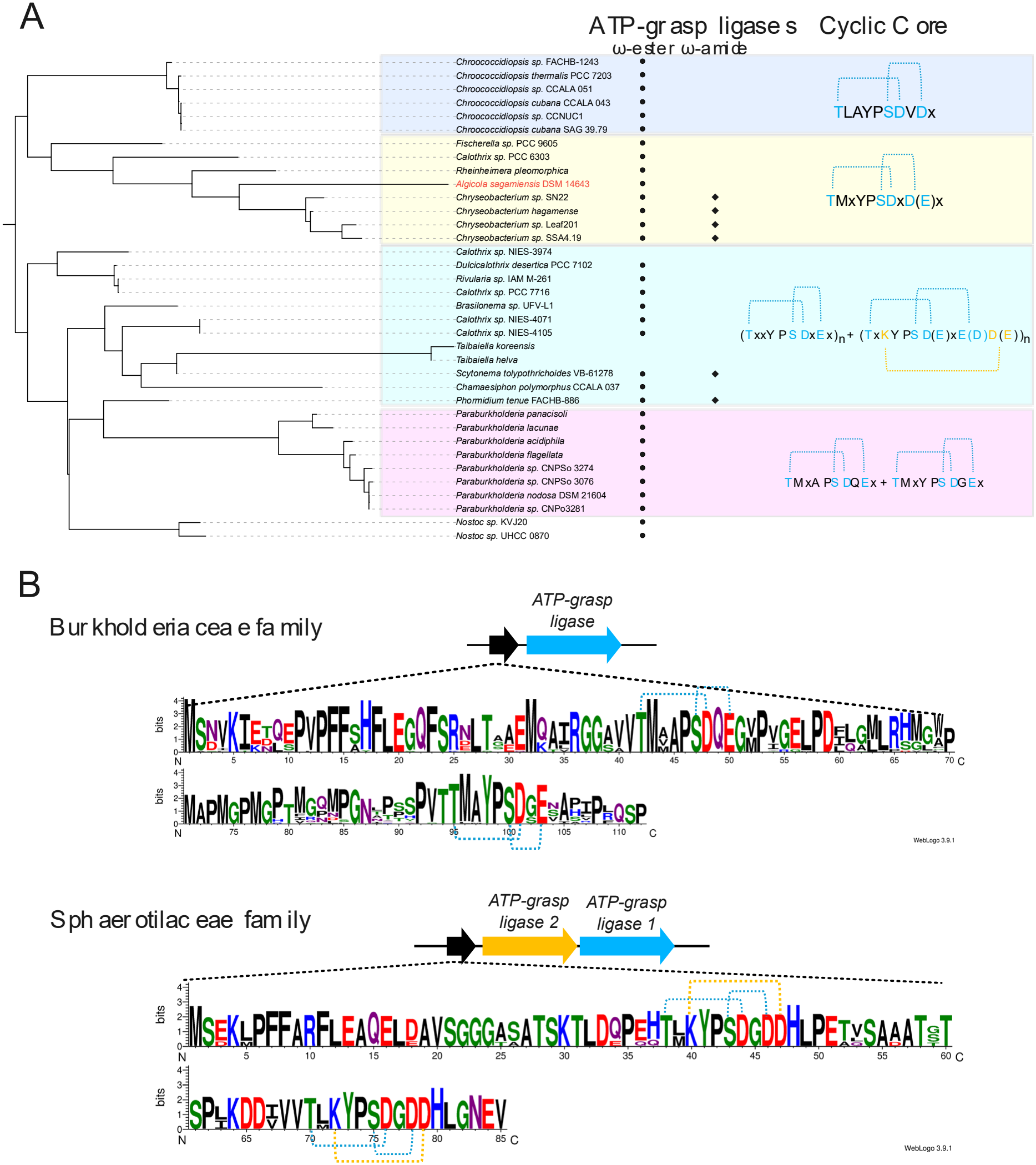
(A) Midpoint-rooted maximum-likelihood phylogenetic tree of putative bicyclic microviridin core-containing precursor peptides identified in this study. “*Algicola sagamiensis* DSM 14643” in red color refers to the natural host strain of the original marinostatin. The presence of associated ATP-grasp ligases is indicated by circles (MvdD-like proteins) and diamonds (MvdC-like proteins). Representative core sequence motifs shared by each clade are shown alongside the tree. (B) Sequence logos of microviridin precursor peptides identified from representative Betaproteobacterial genomes using antiSMASH 8.

Examination of the corresponding genomic contexts revealed discordance between precursor architecture and the ATP-grasp ligase complement expected from the canonical microviridin pathway (Figure 2). In *Taibaiella koreensis* and *Taibaiella helva*, the bicyclic microviridin precursor genes were located on contigs without adjacent microviridin ATP-grasp ligase genes, whereas separate contigs from the same genomes contained complete microviridin biosynthetic loci encoding MvdB-, MvdC-, and MvdD-family proteins. This genomic organization raises the possibility that these orphan precursor genes could be modified in trans by ATP-grasp enzymes encoded elsewhere in the genome. Alternatively, the observed separation could reflect genome assembly fragmentation or errors.

Other lineages displayed a different mismatch between predicted core architecture and biosynthetic enzyme content. Precursors from *Calothrix* sp. NIES-4071 and *Dulcicalothrix desertica* PCC 7102 contain both Lys-deficient bicyclic and Lys-containing tricyclic microviridin-like cores, yet only an MvdD-family ATP-grasp ligase was identified in their corresponding genomes (Figure 2). Under the canonical microviridin biosynthetic model, modification of the Lys-containing core would additionally require an MvdC-family enzyme to install the ω-amide linkage. The absence of an identifiable MvdC homolog therefore raises several possibilities, including broader catalytic functionality of the encoded MvdD-like ligase, the presence of a highly divergent and unrecognized amide-forming enzyme, or presence of a Lys that is not modified to form an ω-amide linkage. Collectively, the interspersed distribution of sequences capable of forming bicyclic and tricyclic cores and the observed discordance between precursor sequence and ATP-grasp gene content suggest that microviridins may have lost or gained the MydC-family ligases and Lys capable of forming the ω-amide cyclization multiple times over their evolutionary history. Against this broader evolutionary heterogeneity, a particularly distinct and conserved lineage was formed by precursors from the order Burkholderiales. Eight closely related precursors from *Paraburkholderia* species in our dataset formed a conserved clade, and each encoded two tandem marinostatin-like cores while lacking a canonical Lys-containing microviridin core (Figure 2). The first core generally followed a TM(A/V/M)APSDQE-like motif, whereas the second was highly conserved as TMAYPSDGE. Despite encoding two putative macrocyclization substrates, the corresponding BGCs consistently contained a single MvdD-family ATP-grasp ligase and lacked an MvdC-family lactam-forming ligase. This conserved combination of tandem Lys-deficient cores and a single ester-forming ATP-grasp enzyme strongly supports the assignment of these loci as a distinct lineage of multicore bicyclic microviridin BGCs.

The distribution of this architecture across multiple *Paraburkholderia* species, including *P. panacisoli, P. lacunae, P. acidiphila, P. flagellata, P. nodosa*, and several unclassified *Paraburkholderia* strains, further suggests that the bicyclic architecture is conserved within this lineage rather than representing an isolated loss of the lactam-forming machinery in a single species. Whereas marinostatins and related bicyclic microviridins have previously been associated primarily with other bacterial lineages, no marinostatin-like natural product from Burkholderiales has, to our knowledge, been structurally and functionally characterized. The conserved and previously unexplored *Paraburkholderia* lineage therefore provided an opportunity to examine whether these predicted multicore precursors are converted into mature bicyclic protease inhibitors.

To determine whether the bicyclic architecture observed in the *Paraburkholderia* clade represents a broader lineage-associated feature, we performed a focused genome-mining analysis of 4,209 representative Betaproteobacterial genomes using antiSMASH 8.^28^ This analysis identified 26 putative microviridin BGCs (SI, Table S4), of which 21 were found in members of the Burkholderiaceae. Strikingly, these Burkholderiaceae BGCs displayed a highly conserved architecture characterized by precursor peptides containing two Lys-deficient, bicyclic microviridin cores. In contrast, the five microviridin BGCs identified outside the Burkholderiaceae encoded precursors containing two canonical, Lys-containing tricyclic microviridin cores and possessed a second ATP-grasp ligase consistent with formation of the additional ω-amide linkage. Phylogenetic analysis of the associated ATP-grasp enzymes, using the characterized MvdC and MvdD enzymes from the microviridin K pathway as references, further distinguished the ester- and amide-forming ligase lineages (Figure S1). Together, these results reveal a strong association between Burkholderiaceae and a conserved multicore bicyclic microviridin biosynthetic architecture within the surveyed Betaproteobacterial genomes and prompted us to experimentally investigate this previously uncharacterized lineage (Figure 2).

We selected the BGC from *Paraburkholderia* sp. CNPSo 3076 as a representative for experimental characterization and designated it *bks*. The minimal *bks* locus encodes the precursor peptide BksA and an MvdD-family ATP-grasp ligase, BksD. BksA contains two predicted core regions, TMVAPSDQE and TMAYPSDGE, both of which lack the Lys residue required for the canonical microviridin macrolactam. Based on their sequence architecture and the presence of BksD as the sole microviridin-type ATP-grasp ligase, we hypothesized that BksD installs ω-ester linkages within both BksA cores to generate bicyclic products. We therefore next sought to reconstruct BksD-dependent macrocyclization of BksA and determine the structures of the resulting products.

### Heterologous Reconstitution Reveals BksD-Catalyzed Bicyclization of Both BksA Core Peptides

To experimentally validate the predicted bicyclic microviridin biosynthetic activity of the *bks* locus, the precursor peptide BksA and its associated ATP-grasp ligase BksD were heterologously produced in *Escherichia coli* BL21(DE3) and purified separately (SI, Table S1 & S2). Because of the limited solubility of BksA, the precursor was expressed as an N-terminal MBP fusion and subsequently released by TEV protease cleavage and purified by HPLC. Purified BksA was then incubated with BksD in the presence of ATP and Mg2+, and the reaction was monitored by LC-ESI-HRMS. In addition to unmodified BksA, products exhibiting sequential mass losses of 18, 36, 54, and 72 Da were observed, corresponding to the elimination of one to four H_2_O molecules. The −4H_2_O product is consistent with formation of the four ester linkages predicted for complete bicyclization of the two BksA core regions. Optimization of the reaction conditions increased the abundance of the −4H_2_O product, with a 2 h incubation providing the greatest accumulation of fully modified BksA and minimal amounts of partially modified intermediates. Prolonged incubation resulted in conversion of the −4H_2_O product to a predominantly −3H_2_O species, consistent with hydrolysis of one ester linkage under the reaction conditions (Figure 3, Figure S2.).

**Figure 3.**
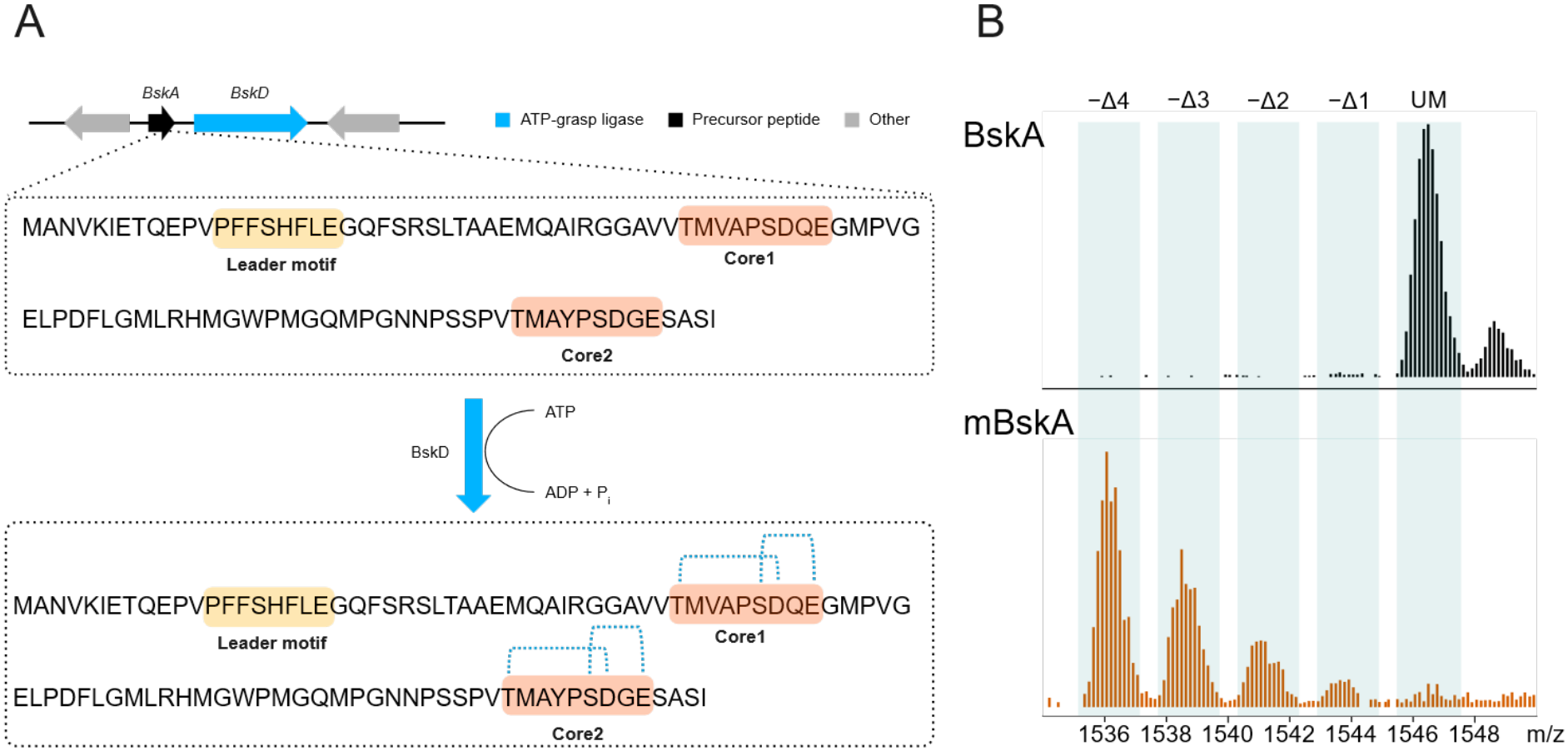
(A) Bicyclic microviridin biosynthetic genes identified from *Paraburkholderia* sp. CNPSo 3076 and proposed macrocyclization of the multicore precursor peptide BksA. (B) LC-ESI-HRMS analysis of unmodified BksA and BksD-modified BksA (mBksA). Peaks corresponding to unmodified precursor (UM) and products exhibiting sequential losses of one to four H_2_O molecules (−Δ1 to −Δ4) are labeled.

To independently assess modification of the two BksA cores and facilitate structural characterization of the mature core peptides, two single-core precursor variants were constructed (SI, Table S1&S2). BksAC1 contained the N-terminal core sequence VVTMVAPSDQEG, whereas BksAC2 contained the C-terminal core sequence PVTMAYPSDGESASI; each retained the BksA leader region required for BksD recognition. Because no obvious dedicated protease could be identified within the *bks* locus and the native proteolytic boundaries of BksA are unknown, an engineered thrombin cleavage site was introduced immediately upstream of each core to enable production of leader-free peptides. Thrombin cleavage leaves an N-terminal GS extension; therefore, the resulting products represent experimentally defined derivatives of the predicted BksA cores rather than established native mature products (Figure 4).

**Figure 4.**
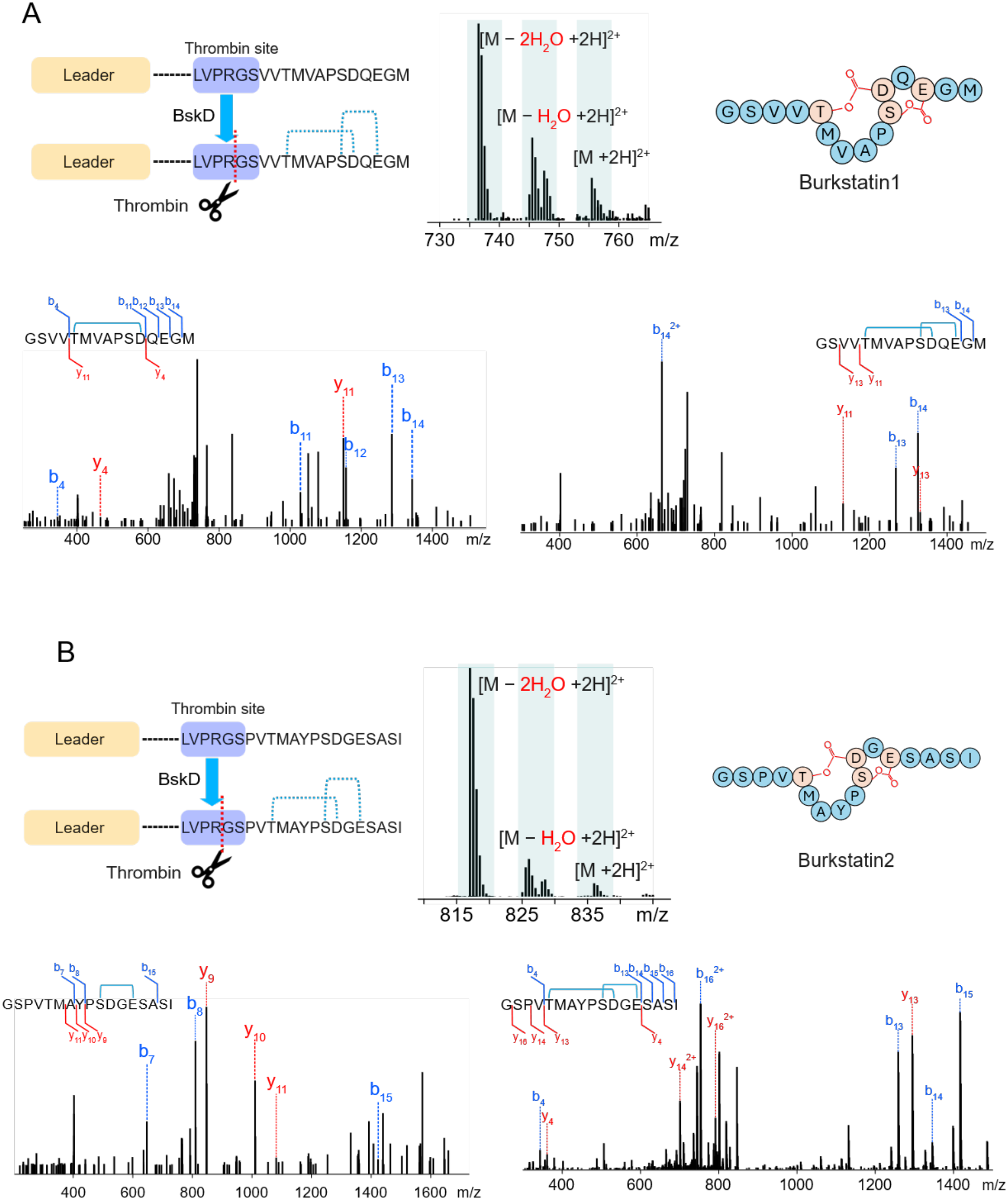
LC-ESI-HRMS and MS/MS characterization of (A) burkstatin 1 and (B) burkstatin Mass spectra of unmodified, monocyclic (−H_2_O), and bicyclic (−2H_2_O) peptides are shown longside the MS/MS fragmentation patterns of monocyclic and bicyclic products. Assigned b- and y-type fragment ions are labeled and mapped onto the corresponding peptide sequences. Proposed ω-ester linkages are indicated in the peptide structures.

Incubation of either BksAC1 or BksAC2 with BksD produced a dominant species exhibiting a 36 Da mass decrease relative to the corresponding unmodified precursor, consistent with two dehydration-dependent cyclization events within each individual core (Figure S2). Together with the fourfold dehydration observed for full-length BksA, these results indicate that a single ATP-grasp ligase, BksD, modifies both BksA cores and installs two macrocycles within each core region. Following BksD modification, the cyclized single-core precursors were treated with thrombin for leader peptide cleavage (Figure S4). The resulting leader-free products were isolated by HPLC for structural characterization. The purified products derived from BksAC1 and BksAC2 were designated burkstatin 1 and burkstatin 2, respectively (Figure 4).

Proposed ω-ester linkages are indicated in the peptide structures.

### LC-MS/MS Analysis Supports the Proposed Ring Topologies of Burkstatin 1 and 2

To further elucidate the ring topologies of burkstatins 1 and 2, both the fully bicyclic products and their corresponding monocyclic intermediates were subjected to LC-MS/MS analysis (Figure 4). For burkstatin 1, MS/MS fragmentation of the bicyclic product revealed two dehydration-dependent modifications localized within the TMVAPSDQE region, consistent with the formation of two ester-linked macrocycles. However, these data alone could not unambiguously distinguish the individual ester linkages. Analysis of the monocyclic intermediate, exhibiting a single dehydration, provided additional structural information. Specifically, b- and y-type fragment ions encompassing the TMVAPSD region retained a mass loss corresponding to one H_2_O molecule, whereas no fragment ions arising from backbone cleavage within the TMVAP region were detected. The absence of fragmentation between Thr and Ser, together with the localization of a single dehydration within TMVAPSD, supported a macrocycle spanning Thr and Asp rather than Ser and Asp. Because Asp is the only acidic residue within this region, these observations supported assignment of the T-D ester linkage in the monocyclic intermediate.

Combined with the two dehydration events localized within TMVAPSDQE in the bicyclic product, these findings supported formation of the second macrocycle through an S–E ester linkage (Figure 4).

For burkstatin 2, LC-MS/MS analysis of the bicyclic product similarly supported two dehydration-dependent cyclizations within the TMAYPSDGE region. In contrast to burkstatin 1, fragmentation of the monocyclic intermediate localized the single dehydration to the SDGE region. Diagnostic b- and y-type ions were observed outside the SDGE region, including fragments separating Thr from the proposed macrocycle, whereas no fragment ions corresponding to backbone cleavage within the SDGE region were detected. This fragmentation pattern supported a macrocycle spanning Ser and Glu rather than Thr and Asp in the monocyclic intermediate. Together with the two dehydration events observed in the bicyclic product, these findings supported assignment of the second macrocycle as a T-D ester linkage (Figure 4).

The proposed ring topologies are also consistent with the established macrocyclization patterns of ω-ester forming ATP-grasp ligases. Characterized MvdD-type enzymes involved in microviridin and marinostatin biosynthesis catalyze ester bond formation between Thr and the first acidic residue (Asp) and between Ser and the second acidic residue (Glu) within their precursor core sequences. Collectively, the complementary LC-MS/MS fragmentation patterns of the monocyclic intermediates and bicyclic products, together with the established biosynthetic precedent, provide strong evidence for the proposed ring topologies of burkstatins 1 and 2.

Although definitive structural confirmation by NMR spectroscopy was precluded by the limited quantities of purified material obtained, the combined mass spectrometric and biosynthetic evidence supports the assigned bicyclic architectures.

### Burkstatins Exhibit Differential Serine Protease Inhibitory Activities

Given the established serine protease inhibitory activities of microviridins and marinostatins, we evaluated the inhibitory activities of burkstatins 1 and 2 against a panel of four serine proteases, including subtilisin, elastase, chymotrypsin, and trypsin. Protease inhibition was assessed using enzyme-specific chromogenic *p*-nitroanilide substrates, which release *p*-nitroaniline upon proteolytic cleavage, allowing enzymatic activity to be monitored spectrophotometrically at 410 nm. Each protease was preincubated with serially diluted burkstatin 1 or 2 at 37 °C for 10 min before addition of the corresponding substrate. Residual protease activity was monitored over 30 min, and concentration-dependent inhibition was evaluated to determine IC_50_ values. All assays were performed in triplicate (Figure 5).

**Figure 5.**
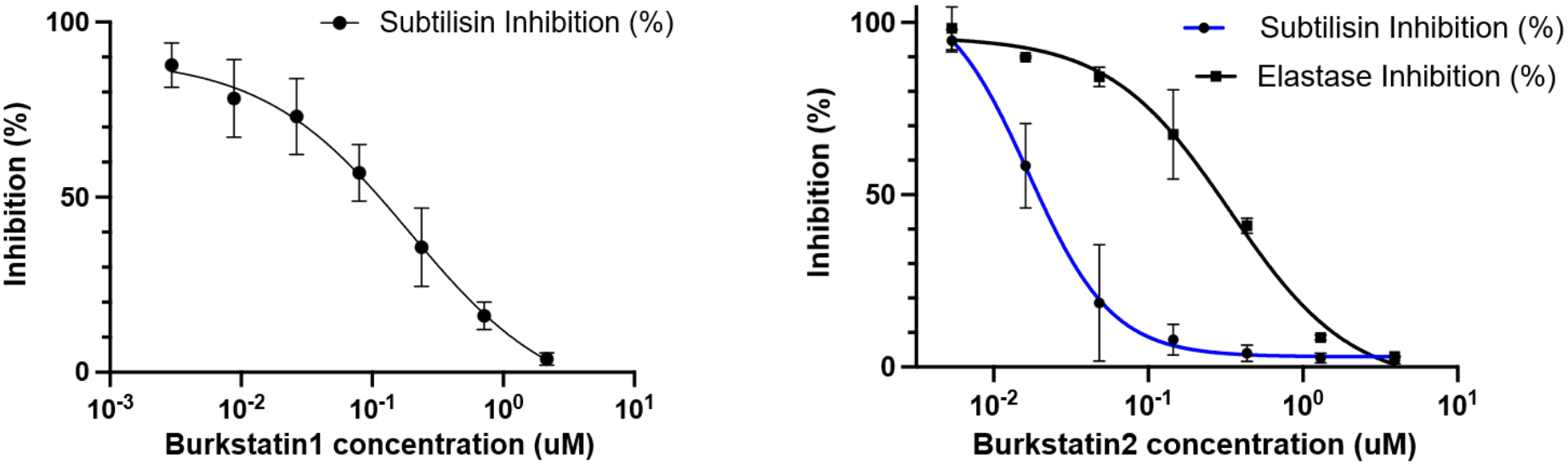
Inhibition of serine proteases by burkstatin 1 and burkstatin 2.

Burkstatin 2 exhibited potent inhibition of subtilisin, with an IC_50_ of 0.020 ± 0.009 μM, and inhibited elastase with an IC_50_ of 0.32 ± 0.07 μM. In contrast, burkstatin 1 displayed weaker inhibition of subtilisin, with an IC_550_ of 0.19 ± 0.09 μM, corresponding to an approximately 9.5-fold difference in potency between the two compounds. No detectable inhibition of elastase was observed for burkstatin 1, and neither compound inhibited trypsin or chymotrypsin under the conditions tested. These inhibitory profiles differ from those of previously characterized marinostatins. Marinostatin has been reported to strongly inhibit chymotrypsin, subtilisin, and elastase, whereas marinostatin E exhibits inhibitory activity against chymotrypsin and subtilisin.^21,26,29^ Marinostatin and marinostatin E differ in the number of amino acids flanking the bicyclic scaffold, indicating that these flanking positions play a role in determining the selectivity of protease inhibition.^21^ Notably, both burkstatins inhibited subtilisin but lacked detectable activity against chymotrypsin, with burkstatin 2 additionally exhibiting inhibition of elastase. The burkstatins differ from each other and from the marinostatins at both the positions flanking the bicyclic core and positions within the bicyclic core. These findings suggest that amino acid sequence variations within and flanking the conserved bicyclic scaffold of marinostatin-type peptides contribute to differences in inhibitory potency and protease selectivity.

## CONCLUSION

In this study, we investigated the biosynthetic diversity of marinostatin-like bicyclic microviridins through genome mining and biochemical characterization. Phylogenetic analysis of microviridin precursor peptides revealed previously underexplored marinostatin-like biosynthetic gene clusters distributed across diverse bacterial taxa, leading to the identification of the *bks* gene cluster from *Paraburkholderia* sp. CNPSo 3076. Heterologous reconstitution demonstrated that the ATP-grasp ligase BksD catalyzes four dehydration-dependent cyclization events across the two core regions of the multicore precursor BksA. Subsequent characterization of the individual core peptides by LC-MS/MS, including analysis of monocyclic intermediates, provided strong evidence for the proposed bicyclic ring topologies featuring Thr–Asp and Ser– Glu ester linkages. The resulting peptides, designated burkstatin 1 and burksatin 2, exhibited different serine protease inhibitory profiles. Together, these findings expand the known diversity of marinostatin-like microviridins, demonstrate the capacity of an ATP-grasp ligase to modify multiple core regions within one precursor, and highlight the influence of core sequence variation on protease inhibitory activity.

## Acknowledgements

Research reported in this publication was supported by the National Institute of General Medical Sciences under award number R35GM146987. The content is solely the responsibility of the authors and does not necessarily represent the official views of the National Institutes of Health. We would also like to thank Vanderbilt’s ACCRE for computational resources.

