## Supplementary material for "Genome-Guided Discovery and Biosynthetic Characterization of Burkstatins, Bicyclic Microviridins with Protease Inhibitory Activity": SI

**Supporting materials and methods**

**Supporting figures**

**Supporting tables**

#### Table of Contents

#### Supporting materials and methods

#### Supporting figures

- Figure S1. Midpoint-rooted maximum-likelihood phylogenetic tree of putative ATP-grasp ligases in betaproteobacterial microviridin BGCs
- Figure S2. LC-ESI-HRMS analysis of unmodified and BskD-modified precursor peptides
- Figure S3. LC-ESI-HRMS analysis of thrombin-digested mBskAC1 and mBskAC2 samples

#### Supporting tables

- Table S1. Primers used in this study
- Table S2. DNA sequences used in this study
- Table S3. Putative bicyclic microviridin precursors identified in this study
- Table S4. Putative microviridin precursors identified in Betaproteobacterial genomes
- Table S5. HR-MS/MS mass spectral data of burkstatins and monocyclic derivatives
- Table S6. HR-MS data of precursor peptides, modified precursors, and burkstatins

### Materials and methods

##### Genome Mining and Phylogenetic Analysis of Microviridin Biosynthetic Gene Clusters

An initial genome-mining survey was conducted to investigate the sequence diversity and phylogenetic distribution of bicyclic microviridin precursor peptides. Microviridin biosynthetic gene clusters (BGCs) were identified using antiSMASH 5, and the corresponding precursor peptide sequences were extracted from the predicted BGCs. Precursors containing bicyclic core architectures were identified based on the absence of residues required for formation of the ω-amide linkage characteristic of tricyclic microviridins. These bicyclic microviridin precursors were subsequently selected for comparative sequence and phylogenetic analyses. Precursor sequences were aligned using MAFFT v7 (L-INS-i), and maximum-likelihood phylogenetic trees were constructed using IQ-TREE 3.

Representative Betaproteobacterial genomes were retrieved from NCBI RefSeq, with one assembly selected per species-level taxon. Non-metagenome-assembled genomes (non-MAGs) were prioritized based on assembly level, completeness, contamination, contig number, and N50. For taxa lacking non-MAG assemblies, metagenome-assembled genomes (MAGs) were included if they exhibited $\geq$ 95% completeness and $\leq$ 5% contamination. In total, 4,209 representative genomes were selected, of which 4,207 were successfully analyzed using antiSMASH 8 with the --fullhmmer, --cb-knownclusters, and --rre options.

Microviridin biosynthetic gene clusters (BGCs) were identified based on antiSMASH predictions and manually inspected. Precursor peptide sequences were extracted from GenBank files using CDS annotations containing "microviridin," together with their corresponding protein identifiers and taxonomic information. Predicted precursor core architectures were manually classified as bicyclic or tricyclic based on sequence motifs. Additional bicyclic microviridin precursor sequences from previously analyzed bacterial genomes were incorporated for comparative analysis.

Precursor peptides and their associated ATP-grasp ligases were independently aligned using MAFFT v7 (L-INS-i). Maximum-likelihood phylogenetic trees were constructed using IQ-TREE 3, with ModelFinder for substitution-model selection, 1,000 ultrafast bootstrap replicates, and 1,000 SH-aLRT replicates. Phylogenetic trees were visualized and annotated using iTOL, with midpoint rooting applied for visualization where appropriate.

##### Construction of expression plasmids

Genes encoding BskD, BskA and single core BskA variants (BskAC1and BskAC2) were synthesized by GenScript. BskD was inserted right after the 6 x His and TEV recognition region of pET-28a by GenScript. BskA, BskAC1 and BskAC2 were inserted right after the TEV protease recognition region of pMAL-c6T.

##### Heterologous expression of BskD, BskA, BskAC1, and BskAC2

Unless stated otherwise, all chemicals were purchased from Research Products International Corp or Sigma-Aldrich. The assembled expression vector pET-28a-*BskD* was co-transformed with a chaperone protein plasmid pGro7 (Takara Bio USA, Inc.) into *E.coli* BL21 (DE3) (New England Biolabs) by electroporation. Transformed cells are cultured on LB plates containing 50 ug/mL Kanamycin and 20 ug/mL Chloramphenicol. Single colonies were picked and cultivated in 5mL LB medium (containing respective antibiotics) overnight at 37°C in a shaker at 250 rpm. The next day, the overnight culture was inoculated into 4 L LB media containing respective antibiotics and 1.5 mg/mL L-arabinose in a shaker at 250 rpm and 37 °C until $OD_{600}$ reached 0.6. IPTG was then added to the cell culture to reach a final concentration of 0.1 mM. The cell culture was then incubated in a shaker at 150 rpm at 18°C for 16 hrs. Expression vectors pMALc6T-*BskAC1 and* pMAL-c6T-*BskAC2* were transformed into *E.coli* BL21 (DE3) by electroporation. Single colonies were picked and cultivated in 5 mL LB medium (containing respective antibiotics) overnight at 37 °C in a shaker at 250 rpm. The next day, the overnight culture was inoculated into 8 L LB media containing respective in a shaker at 250 rpm and 37°C until $OD_{600}$ reached 0.6. IPTG was then added to the cell culture to reach a final concentration of 0.1 mM. The cell culture was then incubated in a shaker at 150 rpm at 15°C for 16 hrs.

**Purification of His-tagged BskD**

After induction, the cells were harvested (4200 rpm, 4°C, 1 hour) and resuspended in lysis buffer (50 mM Tris-HCl, 5 mM imidazole, 300 mM NaCl, 50 mg lysozyme, pH 7.8) and then lysed through sonication. Cell lysate was then centrifuged at 16000 rpm for 20 min at 4 °C. Supernatant was collected and applied to a gravity column with appropriate amount of HisPur Cobalt resin (Thermo Fisher Scientific). After washing out the unbound proteins with 30 column volumes of washing buffer (50 mM Tris-HCl, 20 mM imidazole, 300 mM NaCl, pH 7.8), target protein was eluted using elution buffer (50 mM Tris-HCl, 200 mM imidazole, 300 mM NaCl). Target protein was transferred to a dialysis cassette (Thermo Scientific Slide-A-lyzer Dialysis Cassette, 10K MWCO) and dialysed in 8 L dialysis buffer (50 mM Tris-HCl, 250 mM NaCl, 10% glycerol, pH 7.5) for 12 hours. Protein purity was determined by SDS-PAGE and quantified by BSA assay. Protein was concentrated to ~5 mg/mL by Amicon® Ultra centrifugal filters and then stored at -80 °C for future use.

**Purification of BskA, BskAC1, and BskAC2**

Because BskA, BskAC1, and BskAC2 are all fused to the MBP tag protein with a TEV protease recognition site in between, after heterologous expression, fusion proteins were harvested and crudely purified following the same protocol as purification of His-tagged BskD. After initial purification, fusion proteins were incubated with appropriate amount of TEV protease (TEV protease was expressed and purified in-house using the pET28-MBP-super TEV protease expression plasmid obtained from Addgene; plasmid #171782) (Substrate : TEV = 1: 10) and 10x TEV protease reaction buffer (500 mM Tris-HCl, 5 mM EDTA, 10 mM DTT, pH 7.5) at 4 °C overnight. The protein mixture was then applied to an appropriate amount of HisPur Cobalt resin. After incubation at 4 °C for 2 hours, His-tagged TEV protease and His-tagged MBP proteins were trapped on the cobalt resin and flow-through containing tag free precursor peptides were collected. Precursor peptides were further purified by reversed-phase (RP) HPLC using a C18 column (Hypersil GOLD C18 Selectivity HPLC Column; 100 x 4.6mm, 3μm; Thermo Fisher Scientific). Chromatographic separation was performed at a flow rate of 0.3 mL/min using mobile phase A (95% water, 5% acetonitrile, and 0.05% formic acid) and mobile phase B (95% acetonitrile, 5% water, and 0.05% formic acid). The gradient was maintained at 0% B for 1 min, followed by a linear increase to 100% B over 24 min (1–25 min). BskA eluted at approximately 14.4 min, while BskAC1 and BskAC2 eluted at approximately 13.5 min and 13.3 min respectively. The purified precursor peptides were then lyophilized, validated by LC-MS to confirm molecular weight, and stored at -80°C until use.

**In vitro macrocyclization of BskA, BskAC1, and BskAC2**

Precursor peptides were resuspended with cyclization buffer (100 mM Tris-HCl, 20 mM $MgCl_2$, 5 mM ATP, and 50 mM KCl) and incubated with previously purified BskD (substrate: enzyme = 1:5) at 37 °C on a shaker for 1.5 hrs. For cyclized BskA, cyclization reactions were quenched by adding equal volume of methanol. The mixture was then centrifuged, and supernatant was collected and purified by RP-HPLC using the previously mentioned C18 column and LC set-up.

Modified precursors eluted as the same time as unmodified precursors. Fractions containing cyclized BskA were lyophilized and weighted by analytical balance.

**Thrombin proteolysis releases burkstatin1 and burkstatin2 from leader peptides**

Right after cyclization of BskAC1 and BskC2, thrombin (Sigma Aldrich) was added to cyclization mixture with NaCl and $CaCl_2$ to reach a final concentration of 150 mM NaCl and 2.5 mM $CaCl_2$. After incubation at 4°C overnight, the burkstatin1 and burkstatin2 were purified through RP-HPLC using the previously mentioned C18 column and LC set-up. Burkstatin 1 eluted at approximately 13.0 min and burkstatin 2 at 11.6 min. Fractions containing burkstatin 1 and burkstatin 2 were lyophilized, weighted by analytical balance, and stocked in -80 °C until use.

**Protease inhibition assay**

Lyophilized burkstatin 1 and burkstatin 2 were reconstituted in storage buffer (100 mM Tris-HCl, 50 mM NaCl) to stock concentrations of 47.22 and 258.2 μM, respectively. Trypsin (Thermo Fisher Scientific), elastase (Thermo Fisher Scientific), and chymotrypsin (bioWORLD) were dissolved in assay buffer containing 50 mM Tris-HCl and 20 mM $CaCl_2$ (pH 8.0) to a concentration of 0.1 mg/mL. Their corresponding chromogenic substrates, Nα-benzoyl-DL-arginine 4-nitroanilide (BAPNA) for trypsin, N-succinyl-Ala-Ala-Ala-p-nitroanilide (Suc-AAA-pNA) for elastase, and N-succinyl-Phe-p-nitroanilide (Suc-Phe-pNA) for chymotrypsin, were prepared separately in the same assay buffer at 0.1 mg/mL. Subtilisin (MedChemExpress) and its chromogenic substrate, N-succinyl-Ala-Ala-Pro-Phe-p-nitroanilide (Suc-AAPF-pNA), were separately prepared in phosphate-buffered saline (PBS), each at 0.1 mg/mL. All protease substrates were purchased from Sigma-Aldrich.

Burkstatin 1 and burkstatin 2 were subjected to threefold serial dilutions. For each assay, 2 μL of the diluted peptide solution was mixed with the corresponding protease solution (0.1 mg/mL) and preincubated at 37 °C for 10 min. Subsequently, 40 μL of the corresponding substrate solution (0.1 mg/mL) was added to initiate the enzymatic reaction. Protease activity was monitored spectrophotometrically by measuring absorbance at 410 nm at 37 °C over 30 min using a 384-well plate. All assays were performed in triplicate. (The concentrations of 0.1 mg/mL refer to the separately prepared protease and substrate solutions **before mixing**, not their final concentrations in the reaction mixture.)

**Figure S1.** Midpoint-rooted maximum likelihood phylogenetic tree of putative ATP-grasp ligases in Betaproteobacterial microviridin BGCs (The pink segment of the tree are predicted MvdD enzymes while blue represents predicted MvdC enzymes, labels colored red are ATP-grasp ligases MvdD and MvdC from microviridin K BGC).

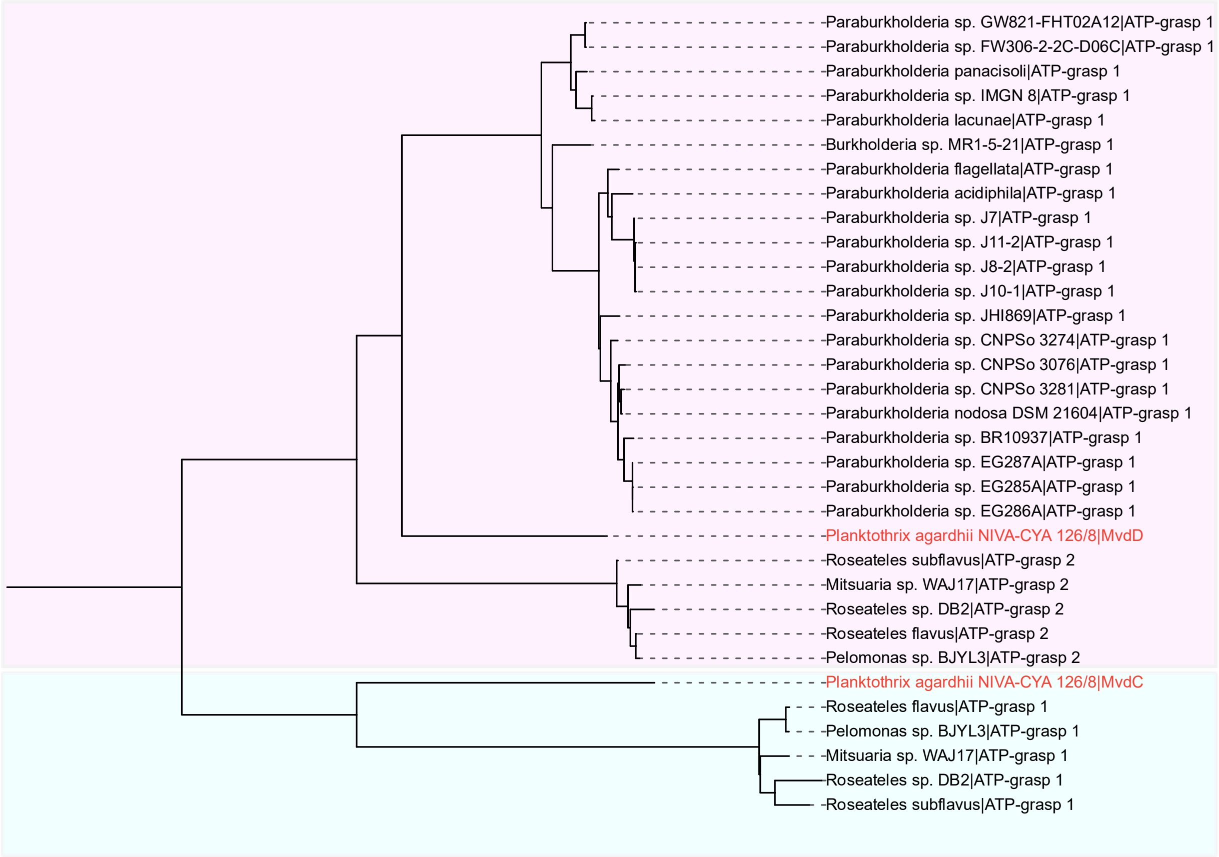

**Figure S2.** LC-ESI-HRMS analysis of unmodified and BksD modified precursor peptides.

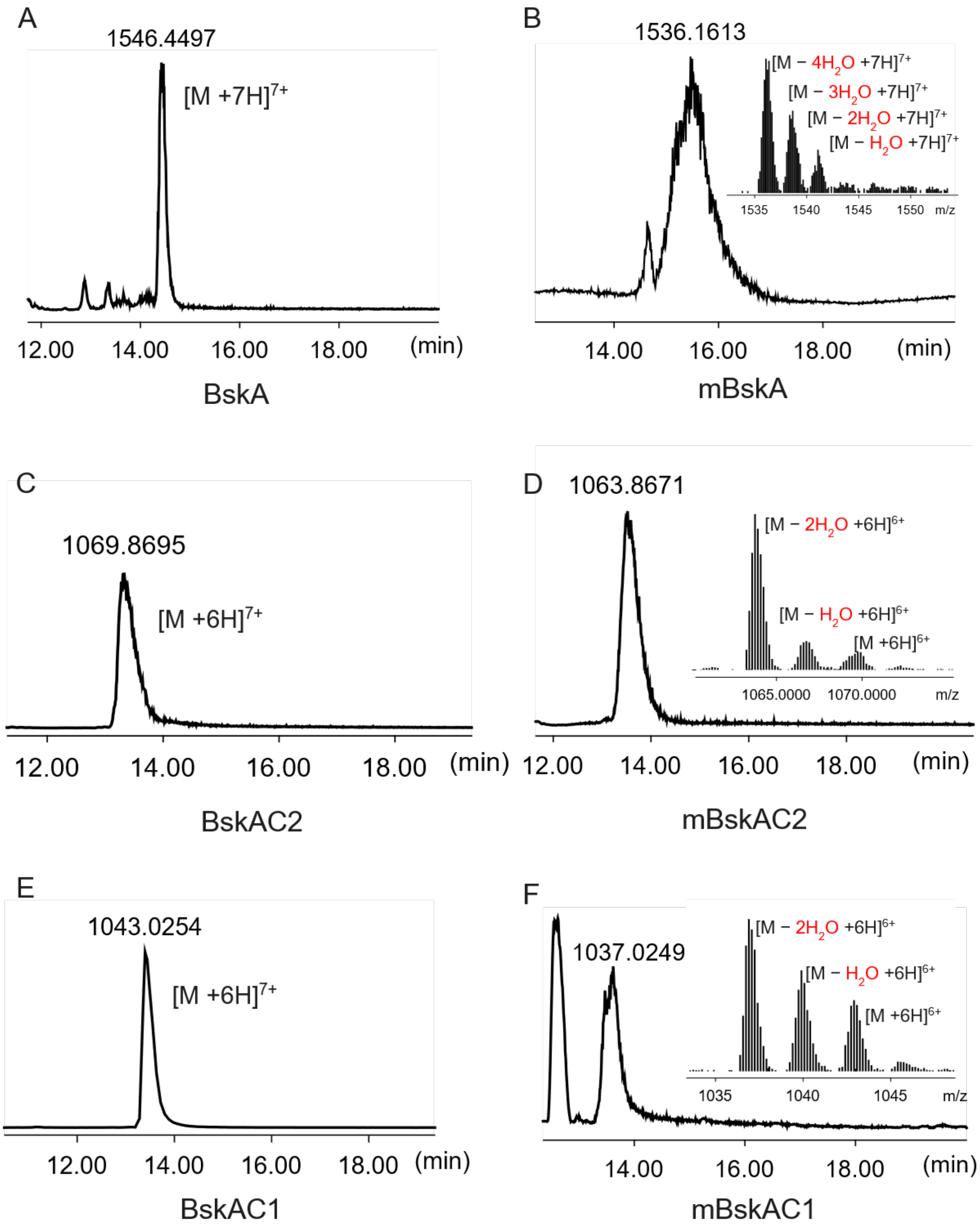

**Figure S3.** LC-ESI-HRMS analysis of thrombin digested mBskAC1 and mBskAC2 samples

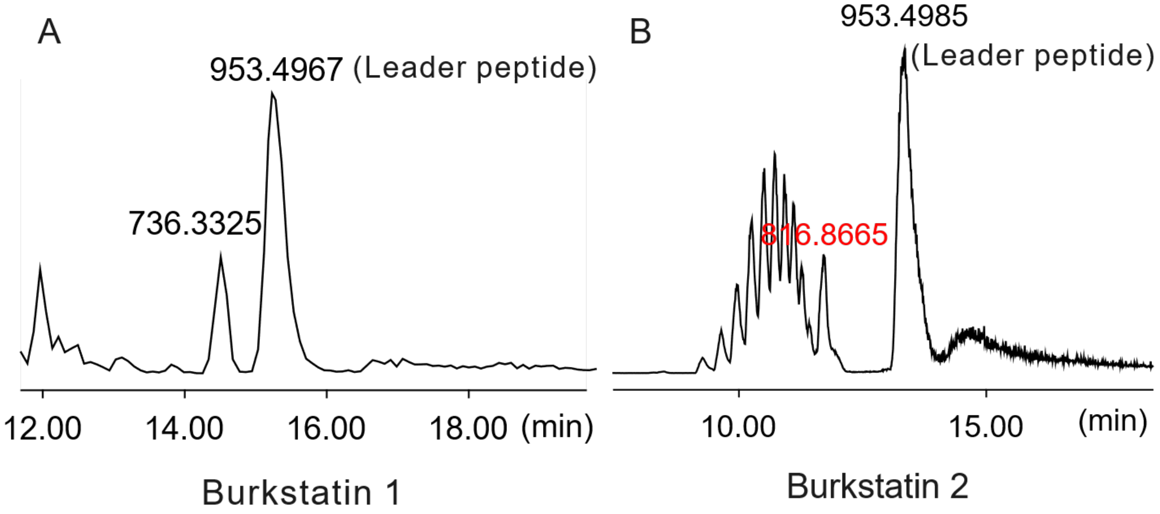

**Table S1**. Primers used in this study

| **Plasmid** | **Primer name** | **Sequence (5' – 3')** |
|---|---|---|
| pMAL-c6T-*BskA* | BskA-F | TAAATGGCACTGCTCGACCTGCTGATGGGCGGCCGCGAT |
| | BskA-R | ATTGATCCGCCTGAACCACCCATCTGGAAGTACAGGTTCTCCCCGA |
| pMAL-c6T-BskAC1 | BskAC1-F | TAAATGGCACTGCTCGACCTGCTGATGGGCGGCCGCGAT |
| | BskAC1-R | ATTGATCCGCCTGAACCACCCATCTGGAAGTACAGGTTCTCCCCGA |
| pMAL-c6T-BskAC2 | BskAC2-F | TAAATGGCACTGCTCGACCTGCTGATGGGCGGCCGCGAT |
| | BskAC2-R | ATTGATCCGCCTGAACCACCCATCTGGAAGTACAGGTTCTCCCCGA |

**Table S2**. DNA sequences in this study

| **Plasmid** | **Sequence name** | **Sequence (5' – 3')** |
|---|---|---|
| pMAL-c6T-*BskA* | BskA | GGTGGTTCAGGCGGATCAATGGCGAATGTTAAAATCGAAACACAGGAGCCAGTTCCATTCTTTTCCCACTTTCTTGAGGGCCAATTTTCCCGCTCCTTAACCGCTGCCGAAATGCAAGCTATTCGTGGCGGTGCAGTCGTCACAATGGTCGCTCCTTCGGACCAAGAAGGTATGCCAGTGGGGGAGCTTCCCGATTTTTAGGCATGTTGCGTCACATGGGATGGCCGATGGGTCAAATGCCGGGCAATAATCCCTCCTCTCCCGTTACAATGGCATATCCCTCCGATGGCGAAAGCGCGTCGATTTAAATGGCACTGCTCGACCTG |
| pMAL-c6T-BskAC1 | BskAC1 | GCGAATGTTAAAATCGAAACACAGGAGCCAGTTCCATTCTTTTCCCACTTTCTTGAGGGCCAATTTTCCCGCTCCTTAACCGCTGCCGAAATGCAAGCTATTCGTGGCGGTGCACTGGTGCCGCGCGGCAGCGTCGTCACAATGGTCGCTCCTTCGGACCAAGAAGGTATGTAA |
| pMAL-c6T-BskAC2 | BskAC2 | GCGAATGTTAAAATCGAAACACAGGAGCCAGTTCCATTCTTTTCCCACTTTCTTGAGGGCCAATTTTCCCGCTCCTTAACCGCTGCCGAAATGCAAGCTATTCGTGGCGGTGCACTGGTGCCGCGCGGCAGCCCCGTTACAATGGCATATCCCTCCGATGGCGAAAGCGCGTCGATTTAA |
| pET-28a-*BskD* | BskD | CATATGTCAGAAAGACACCCCAGGCCAGTAGTTTTGATCATCACGCATTCCAATGATAATGAAAGCATCCCGCTCGTCATGAACGCGG |

| | | |
|---|---|---|
| | | TGGCCGCGCAGGGTGGTATGGCATATCGT<br>TTTGATACCAACCGCTTCCCGACAGATATT<br>CGTTTCAGCAGCCACTACAGCGGTTACGA<br>GGAGCGTCAAGTTATTGCGTGTGAAGATT<br>ACGAGCTCGACCTGGGCACCGTGTCTGCT<br>GTGTGGTATCGTCGTATCGCGACTGGCCA<br>GTGTATTCCGGAGGCTATGGATCGTCAAC<br>TGAGACGTGCGGCTGTGGACCAGAGCCG<br>TGCTACGCTGCATGGCATGATTGCGAGCC<br>TGGACGCATTCCACCTGGATCCGGTTCAC<br>CGTCTGCGTCGCGCGGAAAACAAACAGT<br>TGCAACTGAAGATCGCGCGTGAAGTGGG<br>CCTGACCATTCCGCGTACTCTGATAACGA<br>ACGCCCCTGAGGCGGTTCGTAGCTTCGCC<br>CGTGAGTGCCGTCACGGCATGATCATGAA<br>AACCTTGTCTTCCTTTGCAATTTACGAAG<br>AATGCGAGCAAAAAGTTGTTTTTACCTCC<br>ACCGTTCATGATAAGGACTTGGAGCGCCT<br>GGACGACCTGCGCTACTGCCCGGCAATTT<br>TTCAAGAACAGGTTCCGAAAGCCCTAGA<br>GTTGCGCACCACCATCGTGGGTAATCGCG<br>TGTTCACCGCGTCGATCGATAGCCAGCGT<br>TCGGAAAAGGCGAGCGTCGACTGGCGTC<br>GCGATGGCTTAGAGCTGTTGGAGGCCTGG<br>GAACCGTATGCGCTCCCACCAGAAGTCGA<br>GAACGGTCTCCTGAAGCTGATGCATGCAC<br>TGGGTTTGAATTATGGTGCAGCGGACTTC<br>ATCCTGACCCCGGACGGCCGTCACTGCTT<br>TCTGGAAGTTAACCCGGTAGGCGAATTTT<br>TCTGGCTGGAGAAACACCCGGGTCTGCC<br>GCTGTCTGGTGCGATCGCTGACCTGTTGG<br>TGTCCAACCGCCGGGGTAGCTAACTCGAG |

**Table S3**. Putative bicyclic microviridin precursor identified in this study

| Organism | Genome_accession | Phylum | Precursor sequence | Protein_ID |
|---|---|---|---|---|
| *Chroococcidiopsis* sp. FACHB-1243 | GCF_014696895.1 | Cyanobacteria | MKSQNLNSSIPFFARFLEGQFGAELSETEMASVSGGSEYVTLAYPSDVDVDGG NKGYEFPSIDVPKMPSMPEFPKFPSISDYYKE | WP_199316893.1 |
| *Chroococcidiopsis thermalis* PCC 7203 | GCF_000317125.1 | Cyanobacteria | MNSRNLNSPIPFFAHFLEGQFGAELSEAEMASVSGGNEYATTLAYPSDVDVDG GNKGYEFPSIDVPKMPSMPEFPKFPSISDYYKK | WP_041462301.1 |
| *Chroococcidiopsis* sp. CCALA 051 | GCF_003015105.1 | Cyanobacteria | MSDMNSRNLNSPIPFFAHFLEGQFGAELSEAEMASVSGGSEYATTLAYPSDVDV DGGNKGYEFPSIDVPKMPSMPEFPKFPSISDYYKK | WP_106544128.1 |
| *Chroococcidiopsis cubana* CCALA 043 | GCF_003003835.1 | Cyanobacteria | MNSRNLNSPIPFFAHFLEGQFGAELSEAEMASVSGGSEYAATTLAYPSDVDVDG GNKGYEFPSIDVFKMPSMPEFPKFPSISDYYKK | WP_219888977.1 |
| *Chroococcidiopsis* sp. CCNUC1 | GCF_023546805.1 | Cyanobacteria | MSDMNSRNLNSPIPFFAHFLEGQFGAELSEAEMASVSGGSEYAATTLAYPSDVD VDGGNKGYEFPSIDVFKMPSMPEFPKFPSISDYYKK | WP_106168960.1 |
| *Chroococcidiopsis cubana* SAG 39.79 | GCF_003991895.1 | Cyanobacteria | MSDMNSRNLNSPIPFFAHFLEGQFGAELSEAEMASVSGGSEYAATTLAYPSDVD VDGGNKGYEFPSIDVFKMPSMPEFPKFPSISDYYKK | WP_106168960.1 |
| *Fischerella* sp. PCC 9605 | GCF_000517105.1 | Cyanobacteria | MSNSKKQDLNLSVVPFFARYLEGQLCEDLSSEEMEKITGGSEVVTMRYPSDNE DSNPVPMPPSYPTQPNQPINWNLPSYPHFSIPIPPFCRKRK | WP_063748449.1 |
| *Calothrix* sp. PCC 6303 | GCF_000317435.1 | Cyanobacteria | MSTEKQVLPFFVRYLENQAEDMSIEELNEISGGNGQVVTMAYPSDSDTADGGY STEQPKSPFDMPKFSTNLPFPQFGYGSPYIK | WP_015199858.1 |
| *Rheinheimera pleomorphica* | GCF_011634895.1 | Proteobacteria | MKQPLFATLLETQTDLEQVSAGSLTVTNNLKDQLVTGPSSDGMAVTMAYPSDS DGIDPRVGQDS | WP_166838279.1 |
| *Algicola sagamiensis* DSM 14643 | GCF_000374485.1 | Proteobacteria | MKTTPFFANLLASQTRELTENELEMTAGGTASQQSPVQEVPEQPFFATMRYPSDS DEDGFNFPV | WP_018691846.1 |
| *Chryseobacterium* sp. SN22 | GCF_008329705.1 | Bacteroidota | MENKKSKKPFFASFLEKQIQDPEKVKGGSGAITLPAEDNITSVLKDSVTAPNYD AVTMRYPSDSDAAEDLD | WP_149247165.1 |
| *Chryseobacterium hagamense* | GCF_007991455.1 | Bacteroidota | MENKKTKKPFFASFLEKQIQDPEKIKGGSGTITSPEEDNVTGVLKDNVTAPNYD AVTMRYPSDSDASEDLD | WP_146941447.1 |
| *Chryseobacterium* sp. Leaf201 | GCF_001421435.1 | Bacteroidota | MENKKTKKPFFASFLEKQIQDPEKIKGGTSPLTSVLADDVTDAKKDNVTGVLR DNVTGTSLDAVTLRYPSDSDAAQDLD | WP_056223099.1 |
| *Chryseobacterium* sp. SSA4.19 | GCF_022601875.1 | Bacteroidetes | MENKKSKKPFFASYLEKQIQDPEKVKGGTSPITSVLADEVTDTQKDNVTGVLR DNATSPGLDAVTMRYPSDSDSAQDLD | WP_241612302.1 |
| *Calothrix* sp. NIES-3974 | GCF_002368395.1 | Cyanobacteria | MSEQNHQHPQPPTPFFARYLEAQLQDLSEAEIEAISGGGMAVTLAYPSDQEGVE GGVVTQKYPSDHEDVSDGTITTKKYPSDQEDSGFNIPFPNFPGFNLFSGS | WP_096624416.1 |
| *Dulcicalothrix desertica* PCC 7102 | GCF_007830875.1 | Cyanobacteria | MSNNNNDSINSKPPVPFFARYLEGQLKDLSEEEMAEVVGGSTVTTMAYPSDQE GRDDGMMTTKKYPSDQEDSGFSPIPGVIDFDFQSKFDFPRFPKF | WP_127082103.1 |
| *Rivularia* sp. IAM M-261 | GCF_019973055.1 | Cyanobacteria | MSNNNNDSINSKPPVPFFARYLEGQLEDLSEEEMAEVVGGSTVTTMAYPSDQE GRDDGMMTTKKYPSDQEDSGFGAIPGVIDFDFQSKFDFPRFPKF | WP_223984282.1 |
| *Calothrix* sp. PCC 7716 | GCF_019977735.1 | Cyanobacteria | MSNNNNDSINSKPPVPFFARYLEGQLEDLSEEEMAEVVGGSTVTTMAYPSDQE GRDDGMMTTKKYPSDQEDSGFGAIPGVIDFDFQSKFDFPRFPKF | WP_223984282.1 |
| *Brasilonema* sp. UFV-L1 | GCF_012912105.1 | Cyanobacteria | MSDNNKPALNANVVPFFARYLEGQFCEDLTEEEMQEVKGGIGYATTLAYPSDN ESPDNIAYTLKYPSDEEDGFSQIATKKYPSDNDEYTVTQNYPSNSDDDYSVTVE | WP_169216727.1 |
| *Calothrix* sp. NIES-4071 | GCF_002368455.1 | Cyanobacteria | MSHKDEQALPFFSRFLEGQFCDDLSEEEMNEISGGLKLAVPTQKFPPENPDGIV TTLRYPSDNEDGQPIRPPITKKYPSDHEEYAVTKKYPSEGDDWLATTPIPSYSIE | WP_096691283.1 |
| *Calothrix* sp. NIES-4105 | GCF_002368415.1 | Cyanobacteria | MSHKDEQALPFFSRFLEGQFCDDLSEEEMNEISGGLKLAVPTQKFPPENPDGIV TTLRYPSDNEDGQPIRPPITKKYPSDHEEYAVTKKYPSEGDDWLATTPIPSYSIE | WP_096691283.1 |
| *Taibaiella koreensis* | GCF_003545715.1 | Bacteroidota | MKKQQSGKPFFSLFLESQVKKETLQGGDNPYVTLKHPSDSDEFTTGKYPTDSD EVTMAYPSDSDHEGEPA | WP_118972254.1 |
| *Taibaiella helva* | GCF_003545695.1 | Bacteroidota | MKKQQSGKPFFSLFLESQVKQETLQGGDNPYVTLKHPSDNDEFTTGKYPTDSD EVTMAYPSDSDHDNGELS | WP_118950940.1 |

| | | | | |
|---|---|---|---|---|
| *Scytonema tolypothrichoides* VB-61278 | GCF_000828085.3 | Cyanobacteria | MSDSDKQASNDKAVPFFARYLEGQFCEDLSEEEMENVHGGSTEISDAKGGIVTTLKYPSDNEEGGSVITKRHPSDKDDCGGGISVTRKYPSDNEDAVTLKYPSDGDDDIIAIDTVE | WP_048871252.1 |
| *Chamaesiphon polymorphus* CCALA 037 | GCF_003003845.1 | Cyanobacteria | MKASTIKFSLSTLTLLGLAAVAADPTLATSLQVAENTPSQIQTNDSAGKDLPFFARFLEGQDPEVVDNKVALTESPAIQTQLSNSPRELSKKGIVTTRRYPSDTENTGGGIVTTNRYPSDTDDLTGGTVTTTKYPSDAEDNTGGGIVTTKKYPSDAEDNTGGNLYRRHRR | WP_219892315.1 |
| *Phormidium tenue* FACHB-886 | GCF_014764505.1 | Cyanobacteria | MSEINVNDLNSPAVPFFARYLEGQSVADLSNEEIDAVRGGLKAITMAYPSDQEGDLGVTKRYPSDLGDIAITHKYPSDGDDDIQYLAAPKL | WP_191132225.1 |
| *Paraburkholderia panacisoli* | GCF_008369935.1 | Proteobacteria | MSDVKIKDLPPVPFFAHFLEGQFSRDLTGEEMKALRGGSAVTMAAPSDQEGVPIGELPDIQGLLRSGLAPIRTNLLAPHPVTMAYPSDGEVAPAPFQSP | WP_149676570.1 |
| *Paraburkholderia lacunae* | GCF_003353175.1 | Proteobacteria | MSDVKIKDLSPVPFFAHFLEGQFSRELTGEEMKALRGGSAVTMAAPSDQEGVPIGELPDIQALLRSGLAPVQNNSLTPHPVTMAYPSDGEVAHVPR | WP_115100519.1 |
| *Paraburkholderia acidiphila* | GCF_009789655.1 | Proteobacteria | MSNLKIETQEPVPFFSHFLEGQFSRDLTGAQMQAIRGGAVVTMAAPSDQEGVPVGELPDILGMLRHMGWPMGQLPGTPASPVTTMAYPSDGENAPL | WP_158761792.1 |
| *Paraburkholderia flagellata* | GCF_021390645.1 | Proteobacteria | MSNIKIETQEPVPFFSHFLEGQFSRDLSAAEMQAIRGGAVVTMAAPSDQEGVPVGELPDLLGMLRHMGWPMGPMPGIPSSPVTTMAYPSDGENAPL | WP_233887828.1 |
| *Paraburkholderia* sp. CNPSo 3274 | GCF_024206475.1 | Proteobacteria | MSNVKIETQEPVPFFSHFLEGQFSRNLTTAEMQAIRGGAVVTMMAPSDQEGMPAGELPDFLGMLRHMGFPMGQMPGIPSSPVTTMAYPSDGENASI | WP_254342689.1 |
| *Paraburkholderia* sp. CNPSo 3076 | GCF_026343935.1 | Proteobacteria | MANVKIETQEPVPFFSHFLEGQFSRSLTAAEMQAIRGGAVVTMVAPSDQEGMPVGELPDFLGMLRHMGWPMGQMPGNNPSSPVTMAYPSDGESASI | WP_266229838.1 |
| *Paraburkholderia nodosa* DSM 21604 | CF_000519185.1 | Proteobacteria | MSNVKIETQEPVPFFSHFLEGQFSRNLTAAEMQAIRGGAVVTMVAPSDQEGMPVGELPDFLGMLRHMGWPMGQMPGNNPSSPVTMAYPSDGESASI | WP_035513180.1 |
| *Paraburkholderia* sp. CNPo3281 | GCF_024206515.1 | Proteobacteria | MSNVKIETQEPVPFFSHFLEGQFSRNLTAAEMQAIRGGAVVTMVAPSDQEGMPVGELPDFLGMLRHMGWPMGQMPGNNPSSPVTMAYPSDGESASI | WP_035513180.1 |
| *Nostoc* sp. KVJ20 | GCF_001712795.1 | Cyanobacteria | MPENNQEQEKNQENTPFFERYLEGQESEEVSEELSEEESDSISGGLKFVTRAYPSDREGYKPINVTMRYPSDRE | WP_069068281.1 |
| *Nostoc* sp. UHCC 0870 | GCF_022063185.1 | Cyanobacteria | MSENNQEQENNQENTPFFERYLEGQESEEASEELSEEESDSISGGLKFALTKRYPSDHEGFRTIRTQRYPSF | WP_237988436.1 |

**Table S4**. Putative microviridin precursors in Betaproteobacterial genomes

| Organism | Family | Genome_assembly | Precursor_accesion | Precursor_sequence | ATP_grasp_1_accession | ATP_grasp_2_accession |
|---|---|---|---|---|---|---|
| *Paraburkholderia nodosa* DSM 21604 | Burkholderiaceae | GCF_000519185.1 | WP_035513180.1 | MSNVKIETQEPVPFFSHFLEGQFSRNLTAAEMQAIRGGAVVTMVAPSDQEGMPVGELPDFLGMLRHMGWPMGQMPGNNPSSPVTMAYPSDGESASI | WP_028204601.1 | |
| *Paraburkholderia lacunae* | Burkholderiaceae | GCF_003353175.1 | WP_115100519.1 | MSDVKIKDLSPVPFFAHFLEGQFSRELTGEEMKALRGGSAVTMAAPSDQEGVPIGELPDIQALLRSGLAPVQNNSLTPHPVTMAYPSDGEVAHVPR | WP_208866255.1 | |
| *Paraburkholderia panacisoli* | Burkholderiaceae | GCF_008369935.1 | WP_149676570.1 | MSDVKIKDLPPVPFFAHFLEGQFSRDLTGEEMKALRGGSAVTMAAPSDQEGVPIGELPDIQGLLRSGLAPIRTNLLAPHPVTMAYPSDGEVAPAPFQSP | WP_149676569.1 | |

| | | | | | | |
|---|---|---|---|---|---|---|
| *Paraburkholderia acidiphila* | Burkholderiaceae | GCF_009789655.1 | WP_158761792.1 | MSNLKIETQEPVPFFSHFLEGQFSRDLTGAQMQAIRGGAVVTMAAPSDQEGVPVGELPDILGMLRHMGWPMGQLPGTPASPVTTMAYPSDGENAPL | WP_158761794.1 | |
| *Mitsuaria* sp. WAJ17 | Sphaerotilaceae | GCF_014174515.1 | WP_183105716.1 | MSEKLPFFARFLEAQELDAVSGGGASATSKTLDQPEQTMKYPSDGDDHLPETLSADATSTSPLKDDIVVTLKYPSDGDDHLGNEV | WP_183105717.1 | WP_220800658.1 |
| *Paraburkholderia flagellata* | Burkholderiaceae | GCF_021390645.1 | WP_233887828.1 | MSNIKIETQEPVPFFSHFLEGQFSRDLSAAEMQAIRGGAVVTMAAPSDQEGVPVGELPDLLGMLRHMGWPMGPMPGIPSSPVTTMAYPSDGENAPL | WP_233887829.1 | |
| *Paraburkholderia* sp. CNPSo 3274 | Burkholderiaceae | GCF_024206475.1 | WP_254342689.1 | MSNVKIETQEPVPFFSHFLEGQFSRNLTTAEMQAIRGGAVVTMMAPSDQEGMPAGELPDFLGMLRHMGFPMGQMPGIPSSPVTTMAYPSDGENASI | WP_254342687.1 | |
| *Paraburkholderia* sp. CNPSo 3281 | Burkholderiaceae | GCF_024206515.1 | WP_035513180.1 | MSNVKIETQEPVPFFSHFLEGQFSRNLTAAEMQAIRGGAVVTMVAPSDQEGMPVGELPDFLGMLRHMGWPMGQMPGNNPSSPVTMAYPSDGESASI | WP_254354855.1 | |
| *Paraburkholderia* sp. CNPSo 3076 | Burkholderiaceae | GCF_026343935.1 | WP_266229838.1 | MANVKIETQEPVPFFSHFLEGQFSRSLTAAEMQAIRGGAVVTMVAPSDQEGMPVGELPDFLGMLRHMGWPMGQMPGNNPSSPVTMAYPSDGESASI | WP_266229837.1 | |
| *Pelomonas* sp. BJYL3 | Sphaerotilaceae | GCF_027257975.1 | WP_269630385.1 | MSEKLPFFARFLEAQELDAVSGGGASATSKTLDQPEHTLKYPSDGDDHLPETVSAAATGTSPLKDDIVVTLKYPSDGDDHLGNEV | WP_269630384.1 | WP_269630383.1 |
| *Roseateles subflavus* | Sphaerotilaceae | GCF_030285365.1 | WP_285981663.1 | MSDKLPFFARFLEAQELEAVSGGASATSKTLDEPQHTLKYPSDGDDHLPETQSAAATTTSPIKDDVVVTLKYPSDGDDHLGNEV | WP_285981664.1 | WP_285981665.1 |
| *Paraburkholderia* sp. IMGN_8 | Burkholderiaceae | GCF_038050405.1 | WP_341318064.1 | MSDVKIKDLSPVPFFAHFLEGQFSRELTGEEMKALRGGSAVTMAAPSDQEGVPIGELPDIQALLRSGLAPIQNNSLTPHPVTMAYPSDGEVAHVPR | WP_341318065.1 | |
| *Paraburkholderia* sp. JHI869 | Burkholderiaceae | GCF_038900775.1 | WP_342950898.1 | MSNVKIENQEPVPFFSHFLEGQFSRNLTPAEMQAIRGGAVVTMVAPSDQEGMPVGEMPDIQCLLRHMGLPTMGQMPGNQHAPSTTMAYPSDSEFIPV | WP_342950899.1 | |
| *Roseateles flavus* | Sphaerotilaceae | GCF_039872765.1 | WP_347612734.1 | MSEKLPFFARFLEAQELDAVSGGGASATSKTLDQPEHTLKYPSDGD | WP_347612733.1 | WP_347612731.1 |

| | | | | | | |
|---|---|---|---|---|---|---|
| | | | | DHLPETVSAAATSTSPLKDDIVVT LKYPSDGDDHLGNEV | | |
| *Paraburkholderia* sp. EG286A | Burkholderiaceae | GCF_040965375.1 | WP_324159342.1 | MSNVKIENQEPVPFFSHFLEGQFS RNLTAAEMQAIRGGAVVTMMAP SDQEGMPVGELPDFLGMLRHMG WPMGQMPGAPTSPVTTMAYPSD SENAHI | WP_368707750.1 | |
| *Paraburkholderia* sp. EG287A | Burkholderiaceae | GCF_040965615.1 | WP_324159342.1 | MSNVKIENQEPVPFFSHFLEGQFS RNLTAAEMQAIRGGAVVTMMAP SDQEGMPVGELPDFLGMLRHMG WPMGQMPGAPTSPVTTMAYPSD SENAHI | WP_368707750.1 | |
| *Paraburkholderia* sp. EG285A | Burkholderiaceae | GCF_040965675.1 | WP_324159342.1 | MSNVKIENQEPVPFFSHFLEGQFS RNLTAAEMQAIRGGAVVTMMAP SDQEGMPVGELPDFLGMLRHMG WPMGQMPGAPTSPVTTMAYPSD SENAHI | WP_368707750.1 | |
| *Paraburkholderia* sp. BR10937 | Burkholderiaceae | GCF_040965695.1 | WP_342925434.1 | MSDVKIENQEPVPFFSHFLEGQFS RNLTAAEMQAIRGGAVVTMMAP SDQEGMPVGELPDFLGMLRHMG WPMGQMPGAPTSPVTTMAYPSD SENAHI | WP_368702744.1 | |
| *Roseateles* sp. DB2 | Sphaerotilaceae | GCF_051857535.1 | WP_439440478.1 | MSEKMPFFARFLEAQELDAVSGG GTAATSKTLDQPEHTMKYPSDGD DHLPEAVAATGTSPIKDDVVVTM KYPSDGDDHLGNEV | WP_439440479.1 | WP_439440480.1 |
| *Burkholderia* sp. MR1-5-21 | Burkholderiaceae | GCF_053427715.1 | WP_448087965.1 | MSSIKTEDLSPLPFFAHFLESQFSK DLTREEMKAIRGGSVVTMAAPSD QEGVPTGELPDIQALIRDVTGRVS QLPGIRPQPVTLMYPSDGEIIPL | WP_448087966.1 | |
| *Paraburkholderia* sp. FW306-2-2C-D06C | Burkholderiaceae | GCF_058781505.1 | WP_492596011.1 | MSDVKIKDLSPVPFFAHFLEGQFS RELTGEEMKAVRGGSVVTMVAPS DQEGVPIRGLPDIQALLPSGLAHF RNDPLTPYPVTMAYPSDGEVAAR PL | WP_492596012.1 | |
| *Paraburkholderia* sp. GW821-FHT02A12 | Burkholderiaceae | GCF_059422005.1 | WP_492596011.1 | MSDVKIKDLSPVPFFAHFLEGQFS RELTGEEMKAVRGGSVVTMVAPS DQEGVPIRGLPDIQALLPSGLAHF RNDPLTPYPVTMAYPSDGEVAAR PL | WP_494950812.1 | |
| *Paraburkholderia* sp. J7 | Burkholderiaceae | GCF_905220155.1 | WP_321963291.1 | MSNVKIETQEPVPFFSHFLEGQFS RNLTSAEMQTIRGGAVVTMAAPS DQEGVPVGELPDLLGMLRHMGW PMAPMGPMGPMPGTPSSPVTTM AYPSDGESAPI | WP_321963290.1 | |
| *Paraburkholderia* sp. J10-1 | Burkholderiaceae | GCF_905220165.1 | WP_321945077.1 | MSNVKIETQEPVPFFSHFLEGQFS RNLTSAEMQTIRGGAVVTMAAPS DQEGVPVGELPDLLGMLRHMGW PMAPMGPMGPMGPMPGTPSSPV TTMAYPSDGESAPI | WP_321945076.1 | |

| | | | | | |
|---|---|---|---|---|---|
| *Paraburkholderia* sp. J11-2 | Burkholderiaceae | GCF_905220185.1 | WP_321913347.1 | MSNVKIETQEPVPFFSYFLEGQFSRNLTSAEMQAIRGGAVVTMAAPSDQEGVPVGELPDLLGMLRHMGWPMAPMGPMGPMGPMPGTPSSPVTTMAYPSDGESAPI | WP_321913348.1 |
| *Paraburkholderia* sp. J8-2 | Burkholderiaceae | GCF_905220195.1 | WP_321937034.1 | MSNVKIETQEPVPFFSHFLEGQFSRNLTSAEMQAIRGGAVVTMAAPSDQEGVPVGELPDLLGMLRHMGWPMGPMGPMGPMPGTPSSPVTTMAYPSDGESAPI | WP_321937036.1 |

**Table S5.** HR-MS/MS mass spectrum data

| Peptide | Calculated (Da) mass | Observed (Da) mass | Error/ppm |
|---|---|---|---|
| Burkstatin 1 - b13 | 1265.583063 | 1265.58 | 2.420228343 |
| Burkstatin 1 - b14 | 1322.604527 | 1322.6012 | 2.515491163 |
| Burkstatin 1 - y11 | 1129.465257 | 1129.4628 | 2.175365718 |
| Burkstatin 1 - y12 | 1228.53367 | 1228.5305 | 2.580311861 |
| Burkstatin 1 - y13 | 1327.602084 | 1327.5986 | 2.624280303 |
| Burkstatin 2 - b4 | 341.181946 | 341.1815 | 1.3072204 |
| Burkstatin 2 - b13 | 1256.525214 | 1256.525 | 0.170310948 |
| Burkstatin 2 - b14 | 1343.557243 | 1343.5568 | 0.329721716 |
| Burkstatin 2 - b15 | 1414.594356 | 1414.5936 | 0.534428825 |
| Burkstatin 2 - b16 (2+) | 751.3168305 | 751.3161 | 0.972292873 |
| Burkstatin 2 - y4 | 377.203076 | 377.2028 | 0.73170135 |
| Burkstatin 2 - y13 | 1292.546344 | 1292.5459 | 0.343507992 |
| Burkstatin 2 - y14 (2+) | 696.311017 | 696.264 | 67.52298736 |
| Burkstatin 2 - y16 (2+) | 788.353413 | 788.3826 | 37.02273564 |
| Monocyclic burkstatin 1 -TD - b4 | 343.197597 | 343.1966 | 2.905031995 |
| Monocyclic burkstatin 1 -TD - b11 | 1026.492458 | 1026.4896 | 2.784238674 |
| Monocyclic burkstatin 1 -TD - b12 | 1154.551035 | 1154.5482 | 2.455499942 |
| Monocyclic burkstatin 1 -TD - b13 | 1283.593628 | 1283.5908 | 2.203189497 |
| Monocyclic burkstatin 1 -TD - b14 | 1340.615092 | 1340.6119 | 2.380996618 |
| Monocyclic burkstatin 1 -TD - y4 | 464.18096 | 464.1787 | 4.868790827 |
| Monocyclic burkstatin 1 -TD - y11 | 1147.475822 | 1147.4727 | 2.720754494 |
| Monocyclic burkstatin 2 -SE - b7 | 644.307223 | 644.3046 | 4.071039259 |
| Monocyclic burkstatin 2 -SE - b8 | 807.370552 | 807.3673 | 4.027890282 |
| Monocyclic burkstatin 2 -SE – b15 | 1432.60492 | 1432.6012 | 2.596668452 |
| Monocyclic burkstatin 2 -SE - y9 | 844.368303 | 844.3655 | 3.319641429 |
| Monocyclic burkstatin 2 -SE - y10 | 1007.431632 | 1007.4263 | 5.292666848 |
| Monocyclic burkstatin 2 -SE - y11 | 1078.468745 | 1078.4672 | 1.432586718 |

**Table S6.** HR-MS mass spectrum data

| Peptide | Charge | Calculated (Da) mass | Observed (Da) mass | Error/ppm |
|---|---|---|---|---|
| BskA | +7 | 1546.462725 | 1546.4497 | 8.422446781 |
| BskA - 1D | +7 | 1543.889114 | 1543.8723 | 10.89040092 |
| BskA - 2D | +7 | 1541.315502 | 1541.3052 | 6.683993539 |
| BskA - 3D | +7 | 1538.741891 | 1538.7345 | 4.803089024 |
| BskA - 4D | +7 | 1536.168279 | 1536.1613 | 4.543308053 |
| BskAC1 | +6 | 1043.023576 | 1043.0254 | 1.748762005 |
| BskAC1 -1D | +6 | 1040.021029 | 1040.0256 | 4.39478293 |
| BskAC1 -2D | +6 | 1037.018483 | 1037.0249 | 6.188253575 |
| BskAC2 | +6 | 1069.872106 | 1069.8695 | 2.435805163 |
| BskAC2 -1D | +6 | 1066.869559 | 1066.868 | 1.461596987 |
| BskAC2 -2D | +6 | 1063.867013 | 1063.8671 | 0.082090461 |
| Burkstatin1 | +2 | 736.33215 | 736.3317 | 0.611137243 |
| Monocyclic burkstatin1 | +2 | 745.3374325 | 745.337 | 0.580274089 |
| Burkstatin2 | +2 | 816.8648685 | 816.8658 | 1.14033549 |
| Monocyclic burstatin2 | +2 | 825.870151 | 825.8688 | 1.63585038 |
| LP from BskAC1&2 | +5 | 953.492816 | 953.4985 | 5.961240509 |
